# Cryo-EM structure of an elusive pre-transport intermediate of the multidrug ABC transporter BmrCD reveals modes of asymmetric drug binding

**DOI:** 10.1101/2021.03.13.435271

**Authors:** Tarjani M. Thaker, Smriti Mishra, Wenchang Zhou, Jose D. Faraldo-Gomez, Hassane S. Mchaourab, Thomas M. Tomasiak

**Affiliations:** Department of Chemistry and Biochemistry, University of Arizona, 1041 E Lowell St. Tucson, AZ 85719, USA; Department of Molecular Physiology and Biophysics, Vanderbilt University, 2215 Garland Avenue, Nashville, TN 37232, USA; Theoretical Molecular Biophysics Laboratory, National Heart, Lung, and Blood Institute, National Institutes of Health, 50 South Drive, Bethesda, MD 20892, USA

**Author notes:** These authors contributed equally to this work (Tarjani M. Thaker, Smriti Mishra).

## Abstract

Vectorial substrate efflux by ATP binding cassette (ABC) transporters, which play a major role in multidrug resistance, entails the ATP-powered interconversion of the transporter between stable intermediates. Despite recent progress in structure elucidation of ABC transporters, a number of such intermediates have yet to be visualized and mechanistically interpreted. Here, we combine single particle cryo-EM, Double Electron Electron Resonance (DEER) spectroscopy with Molecular Dynamics simulations to profile and mechanistically frame the conformation of a hitherto unobserved intermediate in the context of BmrCD, a heterodimeric multidrug ABC exporter from *Bacillus subtilis.* In our cryo-EM structure, BmrCD adopts an inward-facing architecture bound to both ATP and the substrate Hoechst-33342 and is capped by an extracellular domain which undergoes ATP-dependent conformational changes. A striking feature of the structure is a symmetric arrangement of the nucleotide-binding domain (NBD) in the presence of ATP whereas binding of Hoechst at two distinct sites in an acidic pocket stabilizes an asymmetric arrangement of the transmembrane domain architecture (TMD). Mutation of residues coordinating Hoechst in the structure abrogates the cooperative stimulation of ATP hydrolysis. In conjunction with previous studies, our findings suggest a mechanistic role for symmetry mismatch between NBDs and TMDs in the conformational cycle of ABC transporters. Moreover, the resolved structures of bimodally-bound drugs are of notable importance for future rational design and optimization of molecules for targeted transport inhibition of ABC transporters.

**ONE SENTENCE SUMMARY:** Cryo-EM and EPR analysis reveal cooperative substrate binding in BmrCD in an architecture primed for transport.

## INTRODUCTION

ATP binding cassette (ABC) transporters harness the energy of ATP hydrolysis to traffic molecules across lipid membranes. Ubiquitous in all kingdoms of life, ABC exporters efflux a spectrum of substrates, most notably cytotoxic drugs^1–3^. Although the molecular architecture of ABC transporters invariably has two nucleotide binding domains (NBD) with two ATP binding sites (also referred to as the nucleotide binding site (NBS)) and two transmembrane domains (TMDs), a subfamily has evolved a catalytically impaired NBD. Typically assembled as heterodimers, ABC exporters of this subfamily are distinguished by an asymmetric duty cycle of the motor domain where ATP hydrolysis is primarily carried out by the active ATP binding site^1^, also known as the consensus NBS. The mechanistic implications of the asymmetric ATP turnover continue to be an active area of investigation.

Owing to the widespread application of single particle cryo-EM, the structural biology of ABC exporters has witnessed rapid advances in the last few years yielding snapshots of transporters in distinct conformational states. Most notably, a complement of such states was resolved for a bacterial ABC heterodimer, TmrAB^4^. These included inward-facing (IF) conformations, occluded and outward-facing (OF) conformations, and for the first time a post-hydrolysis low energy state where asymmetric binding of ATP and ADP was resolved. Current models of ATP-driven transport posit that ABC transporters rest in an inward-facing open conformation with NBDs slightly separated for high affinity substrate and nucleotide binding, though recent studies report nucleotide binding in this architecture is constitutive at least in exporters^1^. The ATP-substrate-transporter complex is then in a state favorable for transport once both NBDs are aligned to form an intact ATP hydrolysis pocket. Conserved loops (Walker A, Walker B) present in most ATPases, and motifs (Q-loop) that are specific only to ABC transporters coordinate the positioning of a catalytic water by a highly conserved glutamate residue in the Walker B motif resulting in ATP hydrolysis and the subsequent reset of the transport cycle.

Despite the success of the TmrAB investigation in linking intermediates of ATP hydrolysis to distinct conformations, significant mechanistic gaps remain. One critical gap pertains to the sequence of events that occur prior to transition to OF conformations after substrate and ATP binding. Except ABC homodimers, structures of ATP-bound ABC exporters almost invariably have been obtained in a catalytically-impaired background where both NBD catalytic glutamates were replaced with Glutamines^4–6^. Moreover, these structures as well as those of ATP-bound TmrAB, capture outward facing or occluded conformations^5–10^. This has led to a generalized conclusion that ATP binding stabilizes the OF conformation despite evidence that the stability of such a conformation may be transporter dependent^11^. Furthermore, prior to this transition, a loaded IF intermediate bound to substrate and ATP must be postulated. Such an intermediate could be transient for TmrAB, explaining its absence in the cryo-EM maps.

A second, heterodimer-specific mechanistic gap, pertains to the nature of drug binding. Efforts to sensitize drug-resistant cells to various small molecules and chemotherapeutics by inhibiting efflux pumps such as heterodimeric ABC transporters has not yet proven successful^12, 13^. In the absence of a detailed understanding of determinants of transporter/drug interactions, progress on this front remains stalled. Some insights have emerged from recent structures of P-glycoprotein (Pgp) visualizing substrates^14^ and inhibitors^15^ cradled in the vestibule between the TMDs with stoichiometries of one and two per transporter, respectively. In contrast, the peptide substrate was poorly resolved in TmrAB structures precluding an analysis of the role of asymmetry in heterodimer substrate recognition, although substrate binding shifted the equilibrium towards a more open IF conformation^4^.

To address these two central mechanistic gaps, we integrated cryo-EM with Double Electron Electron Resonance (DEER) spectroscopy, Molecular Dynamics (MD) simulations, and biochemical analysis to determine, validate and mechanistically contextualize the structure of a pre-transport, ATP-bound, Hoechst-loaded intermediate of the ABC heterodimer BmrCD from *Bacillus subtilis*. BmrCD was selected on the basis of previous DEER analyses demonstrating that the ATP-bound IF intermediate is relatively stable^11, 16^. We further enriched this intermediate in a catalytically impaired mutant background where putative catalytic residues in both NBDs have been replaced with glutamine. Hoechst is transported by BmrCD in inside-out-vesicles and stimulates ATP turnover of the purified transporter in detergent micelles and nanodiscs^11, 17^. The transporter structure adopts an IF conformation characterized by symmetric, yet disengaged NBDs, but structurally asymmetric transmembrane domains. DEER distributions in the mutant background reveal that the cryo-EM conformation is a minor population in the ensemble. Notably, the cryo-EM maps reveal two antiparallel Hoechst molecules bound in the IF cavity, making contacts primarily with the BmrC protomer and seemingly reflecting the internal symmetry of the protein structure. Microsecond MD trajectories are consistent with this interpretation and reveal the network of polar and aromatic interactions that stabilizes both substrate molecules concurrently. The predicted binding mode and stoichiometry of Hoechst is further interrogated by functional analysis and determination of Hill coefficients in wild-type (WT) BmrCD. We found that Hoechst-mediated stimulation of ATP hydrolysis is cooperative, and mutation of residues in contact with the Hoechst molecules blunts stimulation of ATP hydrolysis. In conjunction with DEER analysis, our structure integrates a number of recent results and fills in a key gap in the mechanistic understanding of ATP-coupled conformational dynamics of ABC exporters.

## RESULTS

### The structure of BmrCD in the substrate and ATP-bound state adopts an inward-open architecture

The structure of BmrCD was determined using single particle cryo-electron microscopy (cryo-EM) (**Fig. 1A-C**; **Table 1**; **Supp. Fig. 1**) of a cysteine-less (C-less) BmrCD bearing glutamine substitution of D500 in BmrC and E592 in BmrD (referred to hereafter as BmrCD-QQ). The transporter was purified into LMNG micelles which were subsequently exchanged for digitonin micelles to facilitate orientation distribution on cryo-EM grids (see methods). BmrCD-QQ for cryo-EM structure determination was pre-saturated with excess ATP and the model substrate Hoecsht-33342 (Hoechst) to stabilize the ligand-bound intermediate. From an initial dataset consisting of nearly 4.3 million particles, we determined a structure to 3.5Å resolution from a subset of 157,021 particles in which the topology of the entire architecture of BmrCD is revealed in sufficient detail to define the topology of a TMD insertion in BmrD, between TM2 and TM3 which we define as the BmrCD extracellular domain (ECD) (**Fig. 1A-D; Supp. Fig. 2**). Notably, this domain forms a cap over the extracellular gate of the heterodimer stabilized in an IF conformation where the extracellular side is closed. Within the TMD we also observe two molecules of Hoechst in a central vestibule and a molecule of ATP in each NBD (**Fig. 1B**). Comparison of the observed geometry to the series of IF structures of the closely related exporter TmrAB highlights a wider IF architecture in BmrCD with NBDs slightly separated (**Fig. 2A**) relative to the narrow inward-facing open (PDB ID: 6raf^4^) (RMSD 5.424 Å) or the inward-facing wide conformation of TmrAB (PDB ID: 6rag^4^; RMSD 3.624 Å), both of which are also stabilized by nucleotide binding in the NBS (**Supp. Fig. 3**, **Supp. Table 2**). In comparison to other prokaryotic exporters of known structure^8–10, 18–21^ and related members of the ABCB^5, 14, 15, 22^ and ABCC^6, 7^ families, BmrCD geometry is most similar in architecture to the heterodimeric exporter TM287/288 asymmetrically bound to AMP-PNP in one NBS (PDB ID: 4q4a^21^; RMSD 3.135 Å) (**Supp. Fig. 4, 5**). However, the conformation of BmrCD is unique in that it is both substrate and ATP-bound yet is inward-facing.

**Figure 1.**
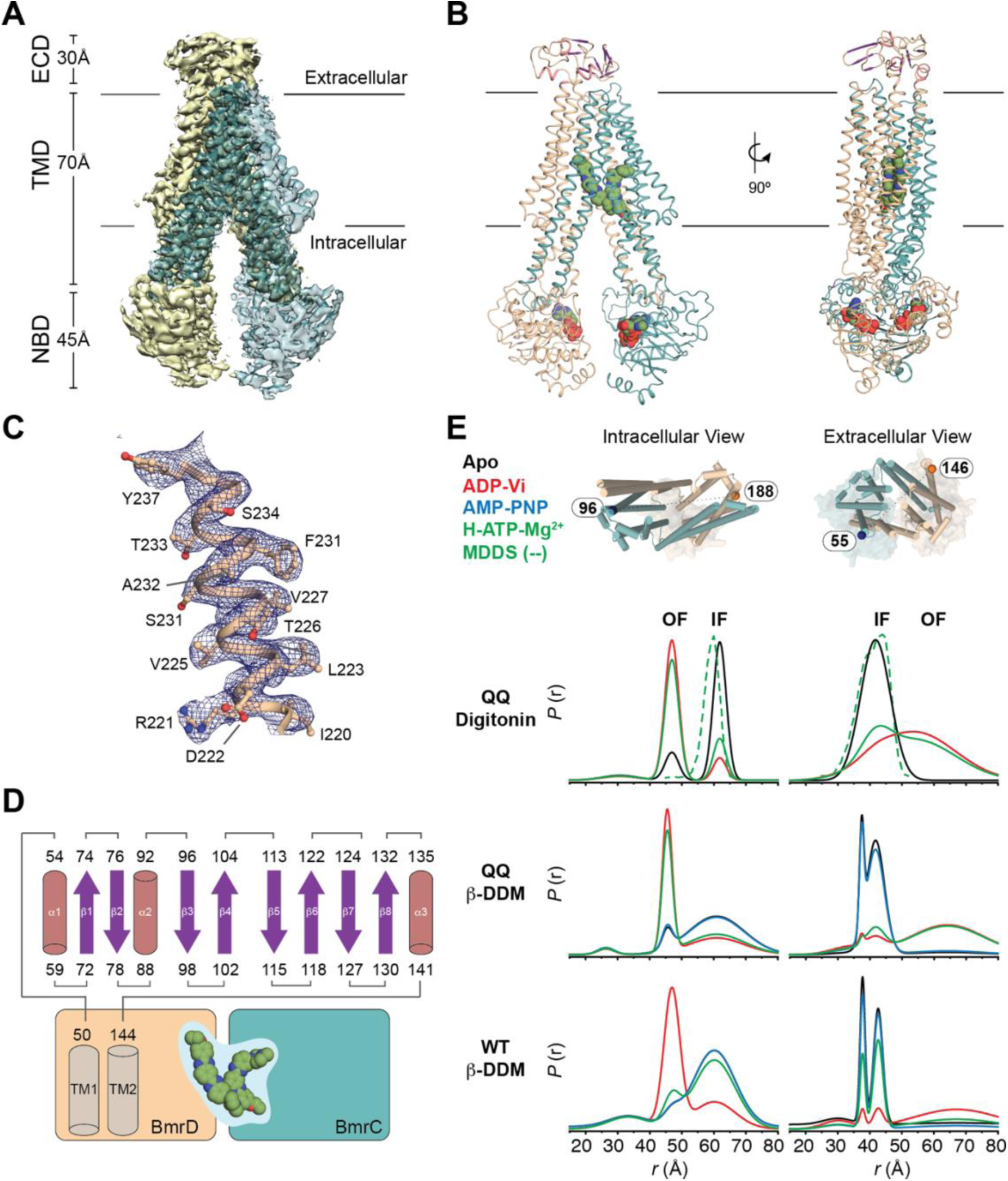
Structure of BmrCD^D500Q/E592Q^ in complex with Hoecsht-33342 and ATP. **A**) Cryo-EM map and **B**) model of BmrCD^D500Q/E592Q^ (BmrCD-QQ) bound to ATP and Hoecsht-33342 (BmrCD-QQ^H/ATP^) highlighting the geometry and arrangement of the transmembrane domains (TMDs) and nucleotide binding domains (NBDs) relative to the extracellular domain insertion (ECD) and the membrane. **C**) Representative density of the 3.5Å resolution structure show in (***A***) and model corresponding to transmembrane helix 3 from BmrD. **D**) Topology of the ECD observed in BmrCD-QQ^H/ATP^ relative to the TMD. **E**) DEER distance measurements for spin label pairs on the intracellular and extracellular regions of the TMD in the cysteine-free wild-type (WT) or the QQ mutant background of BmrCD and in the presence of different substrates. The MDDS-derived distance distributions calculated for the cryo-EM model in (***B***) is overlaid with the DEER derived distance distributions. Peaks corresponding to the inward-facing (IF) or outward-facing (OF) states are labeled as such.

**Figure 2.**
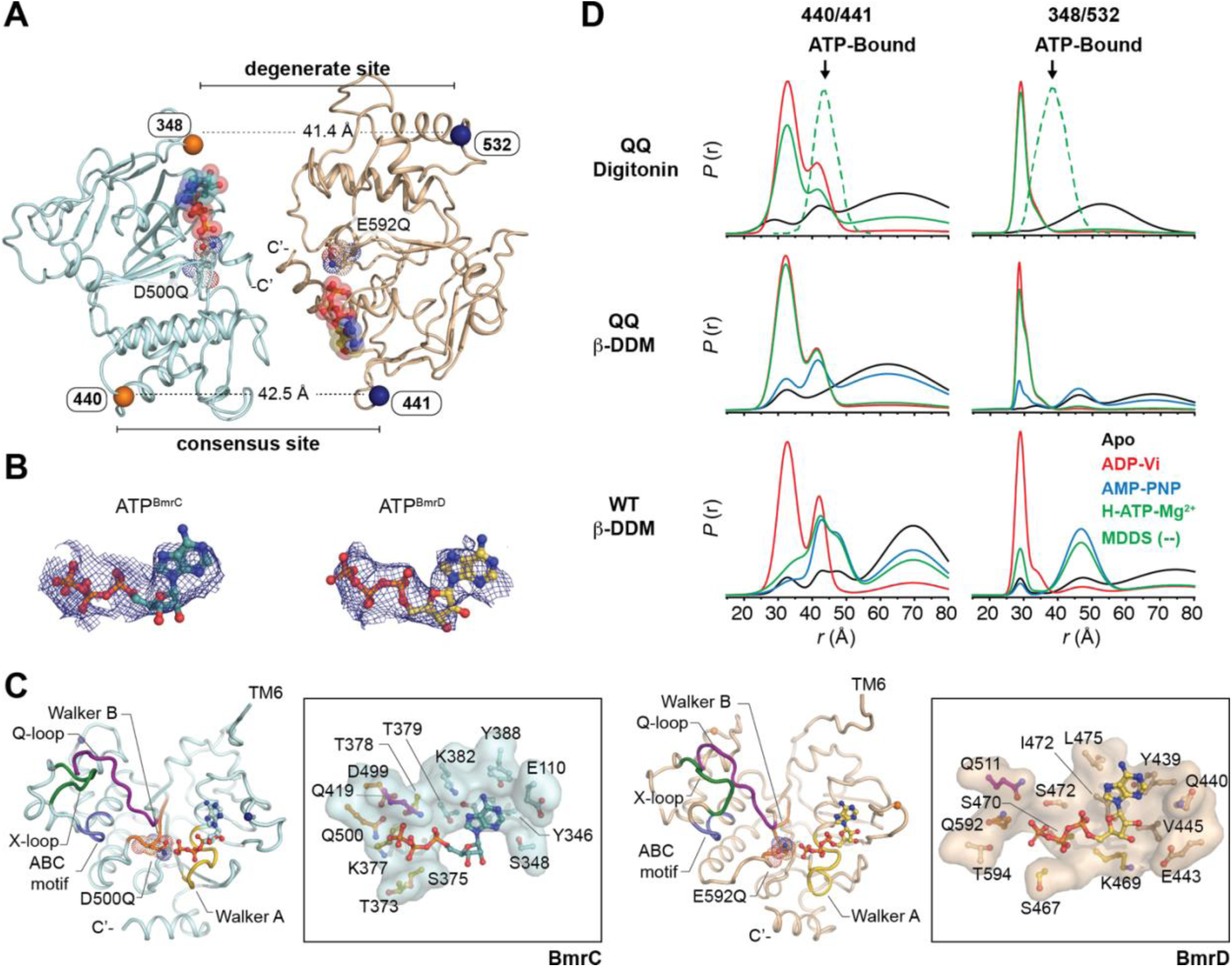
Symmetric geometry of ATP-bound NBSs in BmrCD-QQ^H/ATP^. **A**) Overview of the nucleotide binding domains in the cryo-EM structure of BmrCD-QQ^H/ATP^. **B**) Electron Potential Density corresponding to ATP bound in the BmrC (left) and BmrD (right) chains with symmetric geometries. **C**) Comparison of NBD secondary structure in BmrC (left) and BmrD (right) highlighting the architecture of conserved motifs and the orientations of ATP molecules relative to the position of the catalytic base (D500Q in BmrC; E592Q in BmrD). In the inset, an overview of the atomic features of the ATP binding pockets of each are shown with residues from conserved motifs colored the same. The orientations shown are of the same view in which BmrD was superimposed onto BmrC to highlight the symmetry of the ATP conformation in each. **D**) DEER distance distributions for spin labeled pairs in C-less WT and the QQ mutant of BmrCD consensus (440/441) and degenerate (348/532) nucleotide binding sites under varying conditions. Superimposed in dashed green lines (MDDS) are the distance distributions predicted by the structure.

**Table 1.** Data collection and refinement statistics.

| Refinement |  |  |
| --- | --- | --- |
| Lengths (Å) | 99.55, 150.38, 96.37 |  |
| Atoms | 19430 (Hydrogens: 9679) |  |
| Residues | Protein: 1237 |  |
| Water | 0 |  |
| Resolution (Å) | 3.54 |  |
| Resolution Estimates (Å) | Masked | Unmasked |
| d FSC model (0/0.143/0.5) (Å) | 2.7/3.1/3.7 | 2.9/3.2/6.3 |
| B-factors (Å²) |  |  |
| Iso/Aniso (#) | 9751/0 |  |
| Protein (min/max/mean) | 9.64/107.50/57.52 |  |
| Bonds (RMSD) |  |  |
| Length (Å) (# > 4σ) | 0.007 |  |
| Angles (°) (# > 4σ) | 1.016 |  |
| Validation |  |  |
| MolProbity score | 1.37 |  |
| Clash score | 0.57 |  |
| Ramachandran plot (%) |  |  |
| Outliers | 0 |  |
| Allowed | 6.33 |  |
| Favored | 93.67 |  |
| Rotamer outliers (%) | 2.19 |  |
| Model vs. Data |  |  |
| CC (mask) | 0.69 |  |
| CC (box) | 0.61 |  |
| CC (peaks) | 0.50 |  |
| CC (volume) | 0.65 |  |
| EMRinger Score | 2.57 |  |

### The ATP and Hoechst-bound IF conformation has symmetric but disengaged NBDs

Because previous structures of ATP-bound catalytically impaired ABC transporters have captured occluded or OF conformations^5, 6, 8^, we characterized the ensemble of conformations of the impaired BmrCD-QQ background, solubilized either in digitonin or β-DDM micelles, using DEER distance measurements between pairs of spin labels. Consistent with these previous studies, distance distributions in the TMD (spin label pairs 96/188 and 55/146) report a predominantly OF conformation in the presence of Hoechst and ATP although a minor IF population, which we presume is captured in the cryo-EM structure, is evident in the distance distributions (**Fig. 1D**). Corresponding DEER distributions in the wild-type C-less BmrCD background (WT) previously published are shown for reference^11^.

The structure presented here features symmetric NBDs, and NBSs (**Fig. 2A-C**), consistent with DEER data (**Fig. 2D**). Distance distributions between spin labels introduced in proximity to the degenerate (348/532) and consensus (440/441) ATP binding sites for the QQ background are similar to those obtained in the Vanadate-trapped intermediate of C-less BmrCD (**Fig. 2D**). As previously observed for the EQ double mutants in P-glycoprotein (Pgp)^5^, the D500Q and E592Q mutations present in BmrCD-QQ abrogate the asymmetry of the NBS observed under turnover conditions (solid green traces in **Figure 2D**) in the C-less background. The distance distributions at the NBS pairs predicted from the structure overlap with those experimentally observed in C-less BmrCD in the presence of AMP-PNP (dashed green traces in **Figure 2D**).

In the structure, the geometry of ATP binding is also symmetric and adopts nearly identical conformations (**Fig. 2B,C**). The unambiguous density into which ATP was modeled appears in a canonical site for nucleotide binding that overlaps with ATP in TmrAB^4^ and AMP-PNP in TM288/287^21^. In this orientation, the Walker A motif stabilizes the β−phosphate of ATP, whereas the glutamine residue in place of the catalytic base is oriented in proximity to the γ-phosphate in both protomers (**Fig. 2C**). Unexpectedly, the NBDs of BmrC of BmrD do not contact each other, and nucleotide binding is supported entirely by intra-domain interactions that stabilize ATP within a well-defined cleft. These results support the following two conclusions: 1) The NBD architecture of our structure is more consistent with inward-facing open populations of BmrCD that are stabilized by non-hydrolyzable AMP-PNP and 2) this state is more symmetric in the NBDs than in WT BmrCD in the presence of Hoechst and ATP.

### Architecture and ATP-dependent dynamics of the ECD domain

A unique feature that distinguishes BmrCD from other biochemically characterized ABC exporters, is the presence of the ECD. The Electron Potential Density corresponding to this domain reveals a primarily β-stranded domain consisting of two approximate subdomains of similar topology (**Fig. 1D; 3A**). The ECD is positioned in an orientation that results in extensive interactions with the TMD thereby occluding the extracellular TMD bundle of both BmrC and BmrD. Analysis of the electrostatic distribution over the surface of the ECD and at ECD/TMD interface further reveal both acidic and basic patches that extend into the TMD (**Fig. 3B,C**). An acidic stretch of residues extends from the ECD to the BmrC TMD; whereas, the ECD/TMD interface in BmrD is marked by a basic stretch. Somewhat surprising is the formation of a solvent accessible, acidic cavity in the ECD. Binding of both Hoechst molecules in the BmrCD TMD is observed in a similarly acidic vestibule (see below), suggesting translocation of substrate through the TMD may proceeds via interaction with and rearrangement of the ECD.

**Figure 3.**
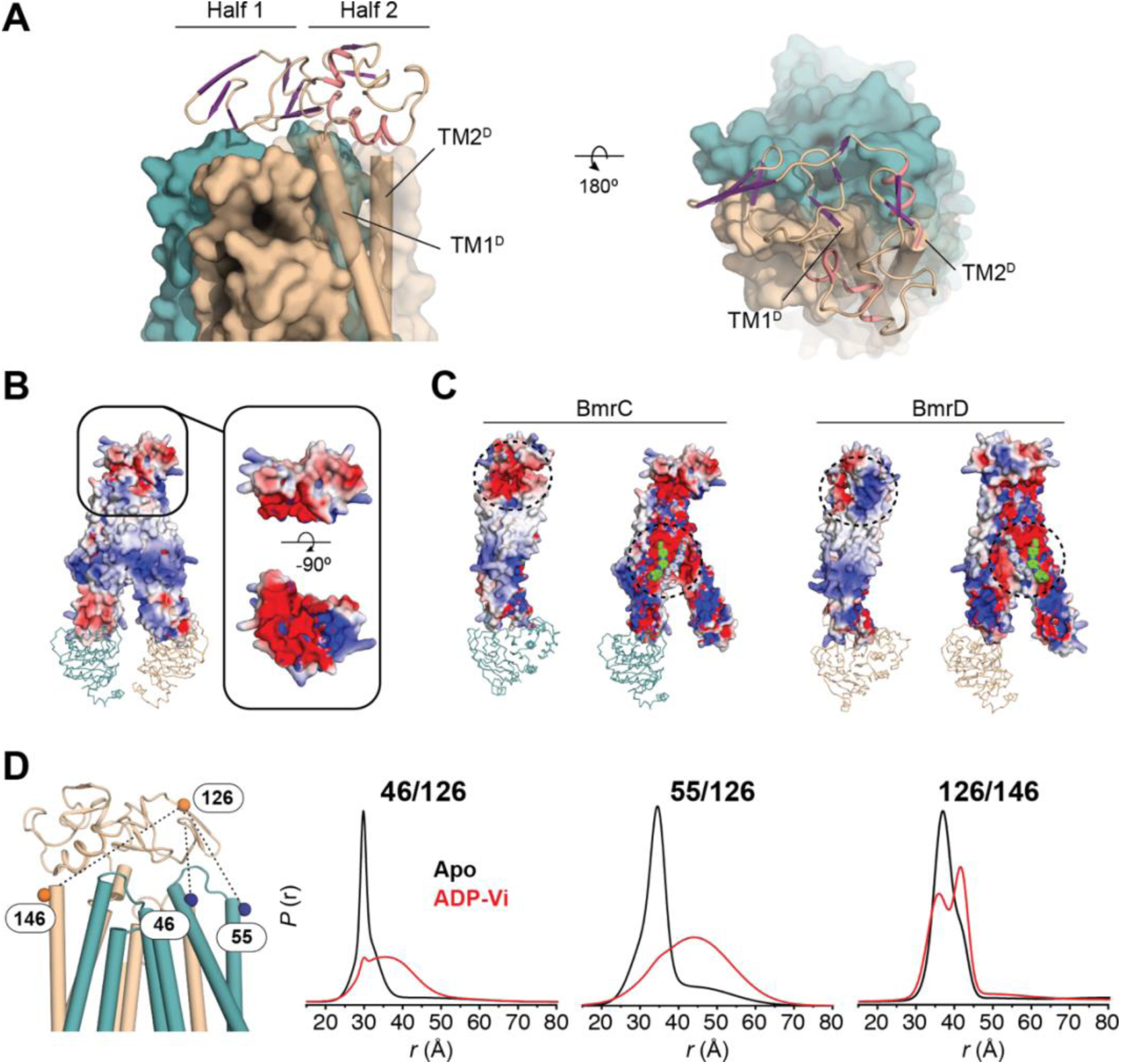
A dynamic role of the ECD in BmrCD catalysis. **A**) Orientation of the ECD in BmrCD-QQ^H/ATP^ relative to its TMD. **B**) Electrostatic distribution over the surface of the BmrCD-QQ^H/ATP^ ECD and its interactions with **C**) BmrC (left) and BmrD (right). **D**) ECD/TMD spin labeled pairs generated in C-less WT BmrCD to monitor the ATP-dependent dynamics of the ECD.

We tested the notion of a functional role for the ECD by measuring distances between a spin label on this domain (BmrD 126) relative to the two spin labels monitoring the extracellular packing of BmrC (55) and BmrD (146) (**Fig. 3D**). The ECD/BmrC interface site (46/126) undergoes a large amplitude distance opening in the high-energy post hydrolysis state (red traces in **Figure 3**). Similarly, the 55/126 pair report a large distance change as would be expected if the ECD moved in concert with BmrD. On the other hand, we observed a small distance change between the BmrD TMD (146) and the ECD (126) suggesting ATP-coupled rearrangement of this domain within the BmrD protomer.

### The ATP and Hoechst-bound IF conformation features asymmetry of the transmembrane domain

Similar to TmrAB and TM287/288, BmrC and BmrD come together to form a vestibule presumably to enable substrate binding. However, this assembly is asymmetric in the TMD of BmrCD (**Supp. Table 1**). Superposition of the BmrD TMD onto the BmrC TMD by rotation around the pseudosymmetric axis of the TMD dimer highlights an outward shift of nearly every TM in BmrC relative to BmrD on the extracellular side (**Fig. 4A**). The arrangement of the intracellular side of the TMD bundle is similarly marked by a concerted outward movement of all BmrC TM helices relative to BmrD. The intracellular side also exhibits the most substantial reorientations, specifically of TM3, 4, and 6 in BmrC which shift away from the substrate binding cavity by ∼4Å relative to BmrD. The presence of several π-helices along the TMD bundle likely accommodates the independent movement of the TMD helices on the two sides of the membrane leaflet.

**Figure 4.**
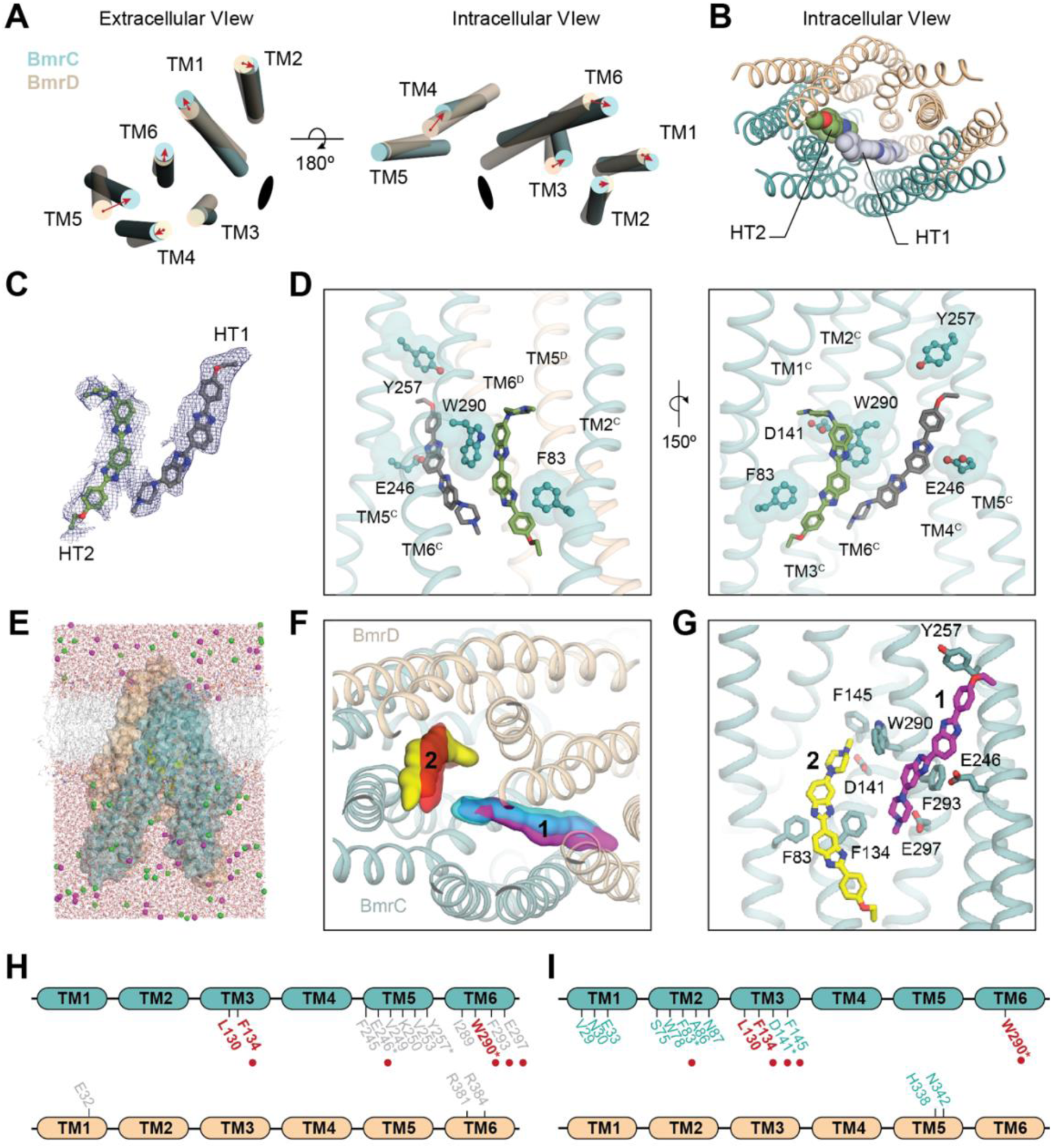
Asymmetric binding of Hoecsht-33342 in BmrCD-QQ^H/ATP^. **A**) Superposition of BmrD transmembrane helices (TM1-6) onto BmrC shown from the extracellular (left) and intracellular (right) gates to highlight asymmetry in the TM arrangement observed in the structure of BmrCD. Red arrows correspond to the movement of each TM in BmrC relative to its position in BmrD. Black ellipse corresponds to the symmetry axis around which BmrD was rotated during superposition. **B**) Intracellular view of the arrangement of two molecules of Hoecsht-33342 (HT1 and HT2) modelled in the TMD cavity of BmrCD. **C**) Electron Potential Density observed corresponding to HT1 and HT2 from a central slice and **D**) their positioning inside of the binding pocket. **E**) All-atom simulation system, comprising the transmembrane domain of BmrCD with two Hoechst molecules placed into the proposed binding site, in a POPC lipid bilayer and 100 mM NaCl (BmrC, *blue* surface/cartoons; BrmD, *orange* surface/cartoons; Hoechst, *yellow* spheres; lipid, *gray* lines; water, *red* lines; sodium, *magenta* spheres; chloride, *green* spheres). **F**) Density maps calculated from the MD trajectories for each Hoechst molecule are overlaid onto the starting cryo-EM structure (*magenta/red* surfaces derive from one trajectory and *cyan/yellow* from the other). **G**) Snapshot representative of the calculated density maps, highlighting the configuration of the two Hoechst molecules and their seemingly most significant sidechain contacts with BmrC. Hydrogen atoms are omitted for clarity. Probability distributions for each of these interaction distances are shown in **Supp. Fig. 6**. The HT1 and HT2 specific interactions are summarized in (***H***) and (***I***), respectively and mapped onto the respective chains colored the same as in (***B***) and (***C***). Residues colored in red are observed interacting with both molecules in the cryo-EM structure. Residues with an asterisk were mutated in this study and are shown as sticks in (***D***). *Red* spheres correspond to residues identified as interacting with HT1 or HT2 in both the cryo-EM structure and all-atom simulation.

Comparison of the pseudosymmetric arrangement of the TMD in BmrCD to the TmrAB TMD highlights similarities to wild-type TmrAB in the nucleotide-bound wide state (PDB ID: 6rag^4^) and the EQ variant in the apo state (PDB ID: 6ran^4^) (**Supp. Table 2**). Even in these related architectures, substantial differences are observed in nearly every TM of BmrD compared to TmrA, the equivalent canonical chain in TmrAB. Comparison of the TMD geometry in related ABC exporters further highlights the asymmetry of the BmrCD TMD halves (**Supp. Fig. 5**). The structures of both apo and AMP-PNP-bound TM287/288 bear the strongest resemblance to the BmrCD TMD geometery^21, 23^. Superposition of each half independently reveals nearly identical bundles in BmrC and TM287, both of which contain the non-canonical NBS and is the site of bound nucleotide in both structures (AMP-PNP in TM287). However, a comparison between the homologous BmrD and TM288 chains reveals a smaller interhelical angle between TM4/TM6 in BmrD (∼50°) than in BmrC or TM287/TM288 (∼60°). These differences could be due to interactions between the BmrD ECD and TMD bundle, or to unique features induced by substrate binding in the BmrCD TMD.

### Asymmetric binding sites of two Hoechst molecules in the TMD

Remarkably, density observed in the TMD vestibule accommodated the assignment of not just one but two molecules of Hoechst. Calculation of a cross-correlation score was used to rationalize a parallel or antiparallel arrangement of the two molecules, with the latter scoring higher (0.65 (antiparallel) versus 0.60 (parallel)) thus supporting an antiparallel assignment reinforced by complimentary local hydrophobic interactions. The Hoechst binding sites are also on each side of a secondary pseudosymmetric two-fold axis relating the NBDs and are positioned asymmetrically relative to the pseudosymmetric axis of the TMD, primarily adjacent to BmrC (**Fig. 4B-D**). Furthermore, binding pocket-lining residues differ between the two halves, i.e. they are non-equivalent from BmrC to BmrD (**Fig. 4H-I**), likely because the TMD segment lining the substrate vestibule itself is asymmetric (**Fig 4A**; **Supp. Fig. 5**). Nonetheless, BmrC residues L130, F134, and W290 form the surface of both binding pockets which consist of chemically similar environment defined by aromatic residues (pocket 1 – F293; pocket 2 – F83 and F134) and capped by an acidic residue (pocket 1 – E297; pocket 2 – D141).

### Antiparallel Hoechst molecules are stabilized by analogous interaction patterns

To evaluate the interpretation of the distinct densities observed within the BmrCD transmembrane domain as two antiparallel Hoechst molecules, and to pinpoint the specific interactions that underpin their stabilization, we turned to all-atom molecular dynamics (MD) simulations (**Fig. 4E-G**). Specifically, we calculated two independent 1-µs trajectories in which the transporter and the two Hoechst molecules are free to diffuse, or reconfigure, and evaluated the resulting structure in each case. For computational efficiency, we considered a protein construct that lacks the nucleotide-binding domains and the ECD inserted between TM1 and TM2 of BmrD was also truncated (**Fig. 4E**). In the cryo-EM structure, these domains contribute minimally to the BmrC-BmrD interface, which primarily involves contacts in the transmembrane domains. The simulated construct showed a high degree of similarity to the starting structure throughout the calculated trajectories, with average backbone RMSD values of 1.6 ± 0.2 Å over the last 0.5 µs (**Fig. 5F**). The two Hoechst molecules initially modeled also remained stably bound, each forming multiple contacts with BmrC, but barely any with BmrD (**Figs. 4G-I; Supp. Fig. 6**). For both molecules, the two simulations resulted in similar poses and interaction patterns; this convergence is reassuring as it indicates both MD trajectories sample the same energetically favorable states. The information from the MD trajectories was used to optimally place the Hoechst molecules in the cryo-EM density.

**Figure 5.**
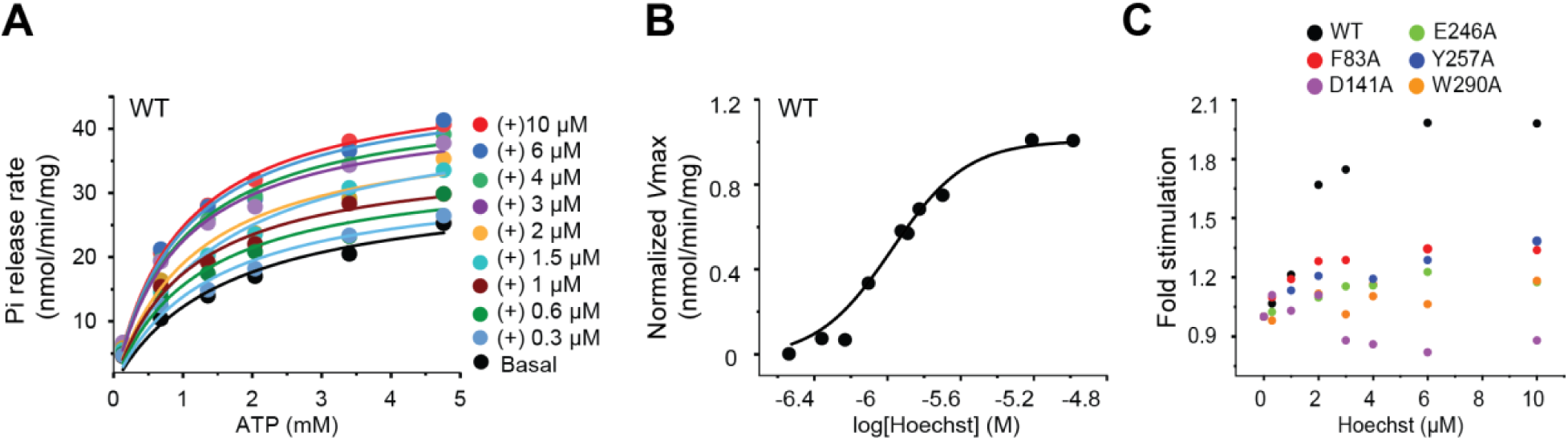
Cooperative stimulation of ATP turnover in BmrCD by Hoechst-33342. **A**) Hoechst-33342 concentration dependent stimulation of ATPase activity in C-less wild-type BmrCD (WT). **B**) Dose-response curve for Hoechst-mediated stimulation of ATPase activity highlighting cooperativity in C-less wild-type BmrCD (WT). The data is a representation of two biological repeats and six technical repeats. **C**) Comparison of Hoechst-stimulated ATPase activity relative to basal activity (fold stimulation) in C-less wild-type (WT) or alanine mutants of BmrCD. Mutants selected are of a subset of residues observed interacting with Hoechst-33342 in the BmrCD-QQ^H/ATP^ cryo-EM structure (see Figure 4).

Unlike the two molecules of ATP in the BmrCD NBSs, the two molecules of Hoechst are marked by different geometries (**Fig. 4C**). The MD simulations show Hoechst-1 (*silver/purple*) is largely coordinated by residues from BmrC TM3, TM5 and TM6 and one residue from BmrD TM1 and two residues from BmrD TM6 (**Fig. 4G,H**) whereas Hoechst-2 (*green/yellow*) forms extensive contacts with residues primarily from BmrC TM2, TM3, and TM6 (**Fig. 4G,I**). The binding pose for Hoechst-1 appears to be defined by the interaction between the piperazine group (carrying a charge of +1) and E297 (**Fig. 4G**). The two benzimidalole rings (which together also carry a charge of +1) form additional π-π and cation-π interactions with the rings of F293 and W290, whereas the etoxy-benzene is engaged by a T-shaped aromatic interaction with Y257. Hoechst-2, while oriented inversely relative to the membrane plane, is similarly anchored by ion-pairing and cation-π interactions formed by the piperazine group, namely with D141, F145 and W290, which is the only residue that contacts both ligand molecules in our trajectories. The benzimidalole rings in molecule ‘2’ form additional π-π and cation-π interactions with the rings of F83 and F134, whereas the etoxy-benzene tail appears to be largely disengaged, in contrast to Hoechst-1 (**Fig. 4G**). These interactions, π-π and cation-π interactions, are reinforced by their observation in the cryo-EM structure. Interestingly, therefore, the pattern of interactions formed by the two molecules in this antiparallel arrangement also appears to reflect the internal structural symmetry of the TMDs.

Lastly, it is worth noting that these interaction patterns recapitulate what is observed in high-resolution crystal structures of other biomolecules in complex with Hoechst. In the outward-open structure of the MFS-family multi-drug transporter LmrP^24^, a Hoechst molecule is stabilized through ion-pairs between two acidic side chains and the piperazine and benzimidalole groups, albeit in an orientation nearly perpendicular to the plane of the transmembrane. The ligand is however only in contact with one of the transmembrane domains, like in our structure of inward-facing BmrCD. Binding of Hoechst to LmrR, a transcriptional repressor of the LmrCD drug ABC transporter, involves aromatic stacking of tryptophan sidechains and the benzimidalole rings^25^, while dipole-π and water-mediated polar interactions explain the mode of binding to DNA^26^. Our cryo-EM and MD simulation data show that it is this interaction versatility that may enable BmrCD to recognize two Hoechst molecules simultaneously.

Moreover, the arrangement and asymmetry in geometry of the two Hoechst molecules in the cryo-EM structure and MD analysis (**Fig. 4C,G**) are not without precedent and are reminiscent of the two molecules of the inhibitor zosuquidar bound to human Pgp^15^ (ABCB1) where one zosuquidar molecule is extended while the other molecule is in a slightly bent conformation (**Supp. Fig. 5**). Binding of these inhibitors occurs at a site closer to the extracellular side, which potentially supports the occluded architecture observed in Pgp. The positioning of Hoechst in BmrCD-QQ is most similar to LPS bound to MsbA and overlaps with the inner core of the polysaccharide portion of the molecule (**Supp. Fig. 5**)^19^, parallel with the presumed inner leaflet of the transmembrane region.

### Cooperative Hoechst stimulation of ATP turnover is blunted by mutation of coordination residues

Because our structure reveals the presence of two bound Hoechst molecules, we reasoned that mutual stabilization would result in cooperative binding of the drug. To test this notion, we determined the k_cat_ of BmrCD ATP hydrolysis as a function of increasing Hoechst concentrations (**Fig. 5A**). We observed a sigmoidal dependence of k_cat_ with a Hill coefficient of approximately 1.8 (**Fig. 5B**). Not only does this data confirm cooperativity, but it also mechanistically validates the binding of two Hoechst molecules by demonstrating direct coupling to ATP hydrolysis in the NBDs.

Conversely, mutations of residues that coordinate the substrate molecules in the structure reduce stimulation of ATP turnover (**Fig. 5C**). We targeted both acidic and hydrophobic residues in the cavity identified by MD and observed in our structure. Alanine substitution effects were variable but overall consistent with the residues implicated in the structure. Notably, W290A which coordinates both Hoechst molecules has a substantial effect on ATP stimulation. Similarly, D141 in proximity to Hoechst-2 abrogates the Hoechst dependence, although its basal rate is higher than for WT BmrCD. This presumably reflects a rearrangement of the BmrC molecules that disrupt coupling of the TMD to the NBD.

## DISCUSSION

The accelerated pace of structure determination of ABC exporters set the stage for an almost unprecedented structure-mechanism understanding. Yet intermediates predicted to be populated in the transport cycle continue to be conspicuously absent from the structural record. In addition, except for TmrAB^4^ and bacterial homodimers^9, 10^, the structure space has been dominated by transporters with impaired ATP-hydrolysis. In the presence of ATP, these models are outward-facing with no evidence of bound substrates. Based on spectroscopic^11, 27^ and biochemical data^28^, the underlying mutations conspire to reshape the energy landscape further confounding the mechanistic interpretation of the structural catalog.

The collection of TmrAB cryo-EM structures is thus far the most complete record of stable intermediates on the energy landscape of an ABC exporter^4^. However, under turnover conditions of excess ATP, two predicted intermediates were missing. One is the outward-facing conformation which purportedly release the substrate, the absence of which was attributed to its presumed transient nature. The structure of the outward-facing conformation was visualized in the catalytically impaired mutant bound to ATP as well as in the vanadate-trapped WT. The second, is a loaded pre-transport intermediate poised for transition to the high energy outward-facing conformation. Because previous structures of catalytically-impaired ABC exporters such as Pgp^5^ and Mrp1^6^ were in the outward-facing conformations, a substrate- and ATP-loaded IF intermediate was not accounted for.

Here, we report the structure of this elusive substrate- and ATP-bound, intermediate. Cryo-EM reveals an IF architecture bound with two molecules of the model substrate Hoechst-33342 in the TMD, and two molecules of ATP in the NBDs of a catalytically impaired variant of BmrCD. Each molecule of Hoechst binds in a similarly located but nonequivalent pocket on the C or D chain and our ATPase data support their role in cooperative allosteric modulation of ATP turnover. Validated by DEER measurements in the WT C-less and the mutant backgrounds, we assign this conformation to the substrate- and ATP-loaded intermediate poised for transition to a high energy, post hydrolysis intermediate. In the case presented here, previous DEER data highlighted the stability of the ATP-bound IF intermediate in BmrCD which was one of two fortuitous factors that allowed it to be represented in our cryo-EM reconstruction, although outward facing states may also be present^11^. The second factor is the structural order of this intermediate which was conducive to it cryo-EM reconstruction.

Our BmrCD structure unlocks details of the nature of pseudosymmetry breaking in ABC transporters and stimulates hypotheses on the mechanistic roles of symmetry mismatch between the NBDs and TMDs. Alignment with the recently determined cryo-EM structures of TmrAB^4^ confirms that BmrCD is in a previously unobserved intermediate. Distances between NBDs are most consistent with the inward-facing wide state, albeit BmrCD is wider than TmrAB. Approximately 75% of BmrCD aligns with this TmrAB conformation, including all of BmrC and most of the transmembrane portion of BmrD. The NBD of the canonical BmrD protomer does not align, instead showing a ∼40° rotation around the symmetry axis of the NBD dimer.

The symmetry of the NBDs holds for most of the transmembrane region but is broken in several locations including TM3 and TM6, especially TM6 in BmrD. TM6 is noteworthy because this helix has been identified as important for substrate gatekeeping in TmrAB^4^ and in *C. elegans* Pgp^29^. In our structure, TM6 in BmrC and BmrD each directly binds one of the two Hoechst molecules which are located in similar pockets in BmrC and BmrD, showing a pseudo-symmetric arrangement. However, TM6 from BmrC breaks that symmetry and is closer to TM3 of BmrC forming a tighter interface than these two helices in BmrD. Moreover, Hoechst binds in a head-to-tail antiparallel arrangement that matches that of inhibitors found in Pgp. MD simulations confirm this arrangement and the stabilizing interactions that largely overlap with the binding mode observed in the structure. We propose a direct relationship between the asymmetry of substrate binding in the cavity and the overall asymmetry of the TMDs of BmrCD.

An emerging theme from structural and spectroscopic investigations of ABC exporters posits a role for substrates and inhibitors in stabilizing symmetric or asymmetric conformations. Cryo-EM structures of Pgp have highlighted different modes of inhibitors and substrates binding with the former filling more of the substrate cavity and inducing a more symmetric arrangement of the NBDs^14^. However, substrate binding induces asymmetric states both in IF conformations and post-hydrolysis conformations, as deduced from DEER analysis of Pgp^30^. Building on this theme, we propose that our symmetry-mismatched BmrCD structure is arrested in an “inhibited-like conformation” by the substitution of the catalytic glutamate in the consensus NBS. In a WT background, we propose that the asymmetry would propagate from the TMD to the NBD resulting in the asymmetric hydrolysis of ATP during transport. Indeed, extensive DEER investigations of BmrCD under turnover conditions concluded that ATP hydrolysis in the consensus NBS is coupled to asymmetric structure of the NBDs^11^. We predict that subsequent or concurrent transition to an OF conformation accompanied by the dissociation of the substrates resolves the asymmetry in the TMD.

In addition to outlining the structural features and the mechanistic context of an intermediate presumed to be poised for transport and revealing modes of substrate interactions with a multidrug transporter, our results have general implications for the field. Our structure resembles a necessary intermediate between symmetric resting states without substrate and asymmetric post-hydrolysis states that have released substrate. In between these states, a transition must occur that “breaks” this symmetry and bring NBDs into alignment for proper catalysis to take place in one of the two NBDs. This transition may reflect asymmetric loading of substrate and asymmetric movements in the TMDs that allosterically facilitate NBD alignment. BmrCD only has one such functional site, but a similar mechanism may be relevant for most ABC exporters, which have two such sites but which nevertheless might have a preferred order^31^. Thus, the absence of intermediates in the structural record of a particular transporter may reflect the energetic idiosyncrasies of the transporter being investigated and should not be extrapolated in the context of a general mechanism. A consensus mechanism for ABC heterodimers will require the convergence of structural, spectroscopic and functional investigations of multiple transporters.

## MATERIALS AND METHODS

### Cloning

Cysteine-less (C-less) BmrCD in pET21b(+) was created as described previously^11^. Briefly, native cysteines in wild-type BmrCD were substituted with alanines using QuikChange site-directed mutagenesis. The C-less BmrCD template was then used to generate BmrCD-QQ where the conserved glutamate of the Walker B motif in the consensus site in BmrD (E592) and aspartate of the degenerate site in BmrC (D500) were substituted with glutamine (Q). Site-directed mutagenesis of the C-less BmrCD was also used to generate double-cysteine pair and substrate binding mutants. All substitutions were confirmed by DNA sequencing.

### Expression and purification

Wild-type C-less BmrCD and all mutant plasmids were transformed into *Escherichia coli* BL21(DE3) cells. A single colony was inoculated into 20 mL Luria Broth (LB) for the primary culture which subsequently was used to start the main culture consisting of 1L of minimal media supplemented with glycerol (0.5% v/v), thiamin (2.5 μg/mL), ampicillin (100 μg/mL), MgSO_4_ (1 mM), and 50X MEM amino acids (1 mL). Cultures were grown at 37°C with shaking to an OD_600_ of 1.2, and the expression of BmrCD induced by the addition of 0.7 mM isopropyl β-D-1-thiogalactopyranoside. BmrCD cultures were incubated at 25°C with shaking for another 5.5 h. The cells were harvested by centrifugation and stored at −80°C. The cell pellets were resuspended in 20 mL of lysis buffer (50 mM Tris-HCl, 5 mM MgCl_2_, 1mM EDTA, pH 7.5), including 10 mM DTT, 10 μg/mL DNase, 0.1 mM PMSF, 1/3 of a Complete EDTA-free protease inhibitor cocktail tablet (Roche) and were lysed by five passes through an Avestin C3 homogenizer at 15-20,000 PSI. The lysate was centrifuged at 9,000g for 10 min to remove cell debris and the membranes isolated by ultracentrifugation at ∼200,000g for 1 h.

Membranes of C-less BmrCD mutants assayed in 0.05% n-dodecyl-β-D-maltopyranoside (β-DDM) were solubilized in resuspension buffer (50 mM Tris-HCl, 100 mM NaCl, 15% (v/v) glycerol, pH 8.0) including 1 mM DTT, 1.25 % w/v β-DDM with constant stirring on ice for 1 h. The membranes of double mutants studied in 0.25% digitonin were solubilized in resuspension buffer (50 mM Tris-HCl, 100 mM NaCl, 15% (v/v) glycerol, pH 8.0) including 1 mM DTT and 1% w/v Lauryl Maltose Neopentyl Glycol (LMNG). Solubilized membranes were then centrifuged at ∼200,000xsg for 1 h to remove insoluble particulates. The solubilized fraction was then incubated for 2 h with 300 µL of pre-equilibrated Ni-NTA resin (QIAGEN) in Ni-NTA buffer (50 mM Tris-HCl, 100 mM NaCl, 15% (v/v) glycerol, 0.05% β-DDM, pH 8.0) for β-DDM samples or Ni-NTA buffer 2 (50 mM Tris-HCl, 100 mM NaCl, 15% (v/v) glycerol, 0.005% LMNG, pH 8.0) for digitonin samples. BmrCD-bound Ni-NTA resin was loaded into a gravity column, washed with five column volumes of Ni-NTA buffer containing 20 mM imidazole and eluted with buffer containing 250 mM imidazole.

### Cryo-EM sample preparation

C-less BmrCD harboring the D500Q (BmrC) and D592Q (BmrD) mutations (BmrCD-QQ) was prepared for cryo-EM by first exchanging sample purified by size exclusion chromatography in SEC Buffer 1 (50 mM Tris-HCl, 150 mM NaCl, 0.01% LMNG, 20% glycerol, pH 7.4) into SEC Buffer 2 (50 mM Tris-HCl, 200 mM NaCl, 0.06% digitonin (Millipore), pH 7.4). Digitonin was added to SEC Buffer 2 from a 10% (w/v) stock prepared by diluting the detergent powder in 50 mM Tris-HCl, 200 mM NaCl, pH 7.4 and boiling at 100°C for 5 min. The solubilized stock was diluted into detergent-free buffer (50 mM Tris-HCl, 200 mM NaCl, pH 7.4) and placed at 4°C for 12-15 hours to allow for impurities in the digitonin to precipitate. Prior to SEC purification, SEC Buffer 2 was filtered through a 0.45 µm filter to remove precipitant from solution. Next, approximately 2.5 mg of SEC-purified BmrCD-QQ was buffer exchanged by two rounds of 10-fold dilution with filtered SEC Buffer 2 and concentration in an Amicon Ultra-100 kDa centrifugal filters (Millipore) prior to SEC purification over a Superose 6 Increase column (Cytiva) equilibrated in SEC Buffer 2 at 4°C (**Supp. Fig. 1**). Fractions were pooled and concentrated to a final concentration of 3.1 mg/mL measured by BCA Assay (Pierce). For substrate trapping, protein was diluted to a final concentration of 22 µM (∼2.5 mg/mL) in 50 mM Tris-HCl, 200 mM NaCl, 0.2% digitonin, pH 7.4 and incubated with 75 µM Hoecsht-33342 (Thermo Fisher) on ice for 30 min prior to the addition of 8 mM ATP/MgCl_2_. To initiate substrate trapping, the sample was heated for 1 min at 37°C following the addition of ATP/MgCl_2_, then briefly placed on ice prior to immediately applying to freshly glow-discharged grids for cryo-preservation.

### Cryo-EM data acquisition and processing

Cryo-EM data were collected on a Quantifoil 1.2/1.3 200-mesh spacing copper grid (Electron Microscopy Sciences) loaded with 5 μL of BmrCD-QQ saturated with Hoecsht-33342/ATP/MgCl_2_ (BmrCD-QQ^H/ATP^) and incubated for 10 s prior to blotting on Whatman 1 paper for 4 s and plunge-frozen in liquid ethane using a GP2 plunge freezer (Leica) equilibrated to 10°C and 85% humidity. In total, 6,919 were recorded by beam-image shift on a 300 kV Titan Krios microscope (Thermo Fisher) equipped with a K3 Summit detector (Gatan) operated in super-resolution mode (Pacific Northwest Center for Cryo-EM) at a nominal magnification of 81,000X, corresponding to a pixel size of 0.54 Å. Dose-fractionated movies were acquired at an electron flux of 0.82 e^−1^/Å^2^ per frame (45 frames total) corresponding to a total dose of 37.2 e^−1^/Å^2^. Images were recorded with a target defocus range of -0.8 to 2.1 µm.

Data were processed in RELION 3.0^32^ and 3.1^33^. Frame-based motion correction, 2X binning, and dose weighting were performed using MotionCor2 to generate an image stack with pixel size of 1.059 Å. Defocus values were estimated from motion corrected, dose weighted images using CTFFIND4.1^34^. Approximately 1,500 particles were manually picked from an initial subset of micrographs and subject to likelihood-based 2D classification to generate templates for automated particle picking. In total, 4,970,928 particles were picked, extracted at a box size of 288 pixels with 4.0 Å/pixel, split into 4 subsets of 1,242,732 particles, and subject to four rounds of 2D classification and particle selection to eliminate bad particles resulting in a final particle set containing 632,978 particles extracted to the full pixel size (1.059 Å) for 3D classification. The best class selected on the basis of highest resolution and visible transmembrane density yielded 157,021 particles, which were then subject to iterative rounds of 3D refinement, CTF refinement and higher order aberration correction, Bayesian polishing, and postprocessing in RELION to yield a final map at 3.5 Å resolution as defined by local resolution calculation in RELION 3.1. Masks were generated manually in Chimera^35^ from RELION and CisTEM^36^ maps. The use of SIDESPLITTER^37^ extensions in RELION 3.1 were used in the reconstruction step in later rounds of refinement with a large mask to counter the effects of observed over-fitting to the detergent micelle and to ensure that all map density was included. Data processing details are shown in **Supplemental Figure 1**.

Model building was performed in Coot^38^ using RELION postprocessed maps with blurring and sharpening as needed generated in CCPEM^39^. A starting structure was created from TmrAB (PDBID: 6rag^4^) truncated to polyalanine. Real space refinement was performed using Phenix^40^ against RELION postprocessed maps. Secondary structure and reference model restraints were used, as was a high nonbonded energy term to ensure proper geometry. The ISOLDE^41^ plug-in to ChimeraX^42^ was used for difficult regions and to ensure correct orientation of H-bonds. Side-chain validation was performed with EMRinger^43^ in Phenix. Placement of the Hoechst molecules was performed manually in Coot and then further refined with MD simulations (see below) and in Phenix. Modelling of the ECD was aided by EVcouplings^44, 45^. All structure figures were made using ChimeraX and PyMol^46^. EM data and atomic coordinates have been deposited in EMDB (EMD-23641) and the Protein Data Bank (PDB ID: 7M33).

### MD simulations

All simulations were carried out with NAMD 2.12^47^ using the CHARMM36 force field^48, 49^ periodic boundary conditions, constant temperature (298 K) and semi-isotropic pressure (1 atm), and an integration time step of 2 fs. Force-field parameters for Hoechst-33342 were those developed in a previous study^24^. Long-range electrostatic interactions were calculated using PME, with a real-space cut-off of 12 Å; van der Waals interactions were computed with a Lennard-Jones potential cut-off at 12 Å with a smooth switching function taking effect at 10 Å. For computational efficiency, the considered in the simulations comprises only the transmembrane domain, i.e. extracellular (BmrD, residue 50-114) and intracellular domains (BmrC, residue 317-574; BmrD, residue 411-665) were truncated. Two molecules of the ligand Hoechst-33342 were included; the initial binding pose was produced by manual-fitting into the cryo-EM density map. The protein-ligand complex was embedded in a pre-equilibrated palmitoyl-oleoyl-phosphatidyl-choline bilayer, in a 100 mM NaCl buffer, using GRIFFIN^47^. Counterions were added to neutralize the net charge of the protein-ligand complex. The resulting system contain 206 POPC lipids and are ∼90 x 90 x 126 Å in size, totaling ∼106,000 atoms. The molecular system was equilibrated following a staged protocol comprising a series of restrained simulations. The protocol consists of both positional and conformational restraints, gradually weakened over 100 ns, and individually applied to protein sidechains and backbone as well as the Hoechst molecules. Subsequently, two trajectories of 1 μs each were calculated, with no restraints.

### Spin-labeling of BmrCD

For EPR spectroscopy, double-Cysteine mutants generated on the C-less background of BmrCD were eluted following Ni-NTA purification and labeled with 20-fold molar excess of 1-oxyl-2,2,5,5-tetramethylpyrroline-3-methyl methanethiosulfonate (Enzo Life Sciences) at room temperature in the dark over a 4 h period, after which protein samples were placed at 4 °C overnight (∼15 h). The labeled protein samples in β-DDM and LMNG were then separated from free label by size-exclusion chromatography on a Superdex 200 Increase column (Cytiva) in buffer containing 50 mM Tris-HCl, 150 mM NaCl, 10% (v/v) glycerol, pH 7.4 containing 0.05% β-DDM (β-DDM samples) or 0.01% LMNG (digitonin samples). The samples in 0.01% LMNG containing buffer were then exchanged into digitonin buffer (50 mM Tris-HCl, 200 mM NaCl, 0.06% w/v digitonin, pH 7.4). The collected fractions of spin-labeled BmrCD mutants were concentrated using an Amicon Ultra-100 kDa centrifugal filters (Millipore), and the final concentration determined by absorbance at 280 nm (Mean extinction coefficient = 68077.5 M^−1^ cm^−1^).

### DEER sample preparation and DEER spectroscopy

Spin-labeled BmrCD mutants were concentrated to 70-100 µM using Amicon Ultra-100 kDa centrifugal filters (Millipore) and incubated with nucleotides or Hoechst-33342. The final concentrations of ATP, AMP-PNP, vanadate, MgCl^2+^, and digitonin were 10 mM, 10 mM, 5 mM, 10 mM, and 0.25% respectively. Samples for DEER analysis were cryoprotected with 24% (v/v) glycerol. Post-hydrolysis (ADP-Vi) and turnover (H-ATP-MgCl^2+^) samples prepared in digitonin buffer were incubated at 37°C for 15 min and 1 min, respectively. β-DDM-solubilized protein samples were trapped in the post-hydrolysis state and with AMP-PNP by incubating at 30°C for 30 min. Turnover samples in β-DDM buffer conditions were incubated at 30 °C for 5min. Post-hydrolysis (ADP-Vi), turnover (H-ATP-MgCl^2+^) and AMP-PNP reactions were stopped by flash freezing in a liquid nitrogen bath.

DEER spectroscopy was performed on an Elexsys E580 EPR spectrometer operating at Q-band frequency (33.9 GHz) equipped with a 40W Amp-Q amplifier (Bruker) with the dead-time free four-pulse sequence at 83 K^50, 51^. The pulse lengths were 10 ns (p/2) and 20 ns (p) for the probe pulses and 40 ns for the pump pulse. The frequency separation was 73 MHz. Raw DEER decays were analyzed as described previously^11^. Briefly, primary DEER decays were analyzed using home-written software operating in the Matlab environment. The software carries out global analysis of the DEER decays obtained under different conditions for the same spin-labeled position. The distance distribution is assumed to consist of a sum of Gaussians, the number and population of which are determined based on a statistical criterion. Distance distributions on the BmrCD-QQ cryo-EM structure were predicted *in silico* using 1 ns molecular-dynamics simulations with dummy spin labels with default parameters using the DEER Spin-Pair Distributor at the CHARMM-GUI website^52, 53^. Experimental Data related to **Figures 1E**, **2D**, and **3D** are reported in **Supplemental Figure 7**.

### ATPase assays

The specific ATPase activities for wild-type (WT) and mutants of C-less BmrCD were determined by an inorganic phosphate assay as previously described^11^ with slight modification. Briefly, BmrCD (20 µg) samples were incubated with increasing concentrations of ATP at 30°C for 30 min under basal conditions (no Hoechst) or presence of different concentrations of Hoechst. The reaction was stopped by adding 1% SDS and the color was developed using a 1:1 solution of ammonium molybdate (2% in 1M HCl) and ascorbic acid (12% in 1M HCl). The absorbance of samples was measured at a wavelength of 850 nm on a BioTek Synergy H4 microplate reader. The amount of phosphate released was determined by comparison to a standard curve generated from inorganic phosphate. The *V*max of WT and mutant C-less BmrCD was derived using the Levenberg-Marquart nonlinear least squares fitting approach in Origin (OriginLab, Inc).

## ACKNOWLEDGEMENTS

We thank Theo Humphries and other support staff at the Pacific Northwest Center for Cryo-EM (PNCC) for assistance with cryo-EM data collection. We thank members of the Tomasiak lab and Derek Claxton from the Mchaourab lab for critical reading of this manuscript. This research was funded by the Division of Intramural Research of the National Heart, Lung and Blood Institute (W.Z. and J.D.F.G), and grants from the National Institute of General Medicine Sciences awarded to T.M. Tomasiak (R00 GM114245) and H.S. Mchaourab (GM128087). Computational resources were in part provided by the NIH HPC facility Biowulf.

## AUTHOR CONTRIBUTIONS

This study was conceptualized by T.M. Tomasiak and H.S. Mchaourab. Experimental design, investigation, and analysis were conducted by T.M. Thaker, S.M, W.Z., and J.D.F.G., H.S. Mchaourab, and T.M. Tomasiak. The initial manuscript was prepared by T.M. Tomasiak, T.M. Thaker, and H.S. Mchaourab, with further editing by S.M. and J.D.F.G. Data presentation and visualization was conducted by T.M. Thaker and S.M. Supervision of research and funding acquisition was carried out by T.M. Tomasiak and H.S. Mchaourab.

---

**Supplementary Figure 1.**
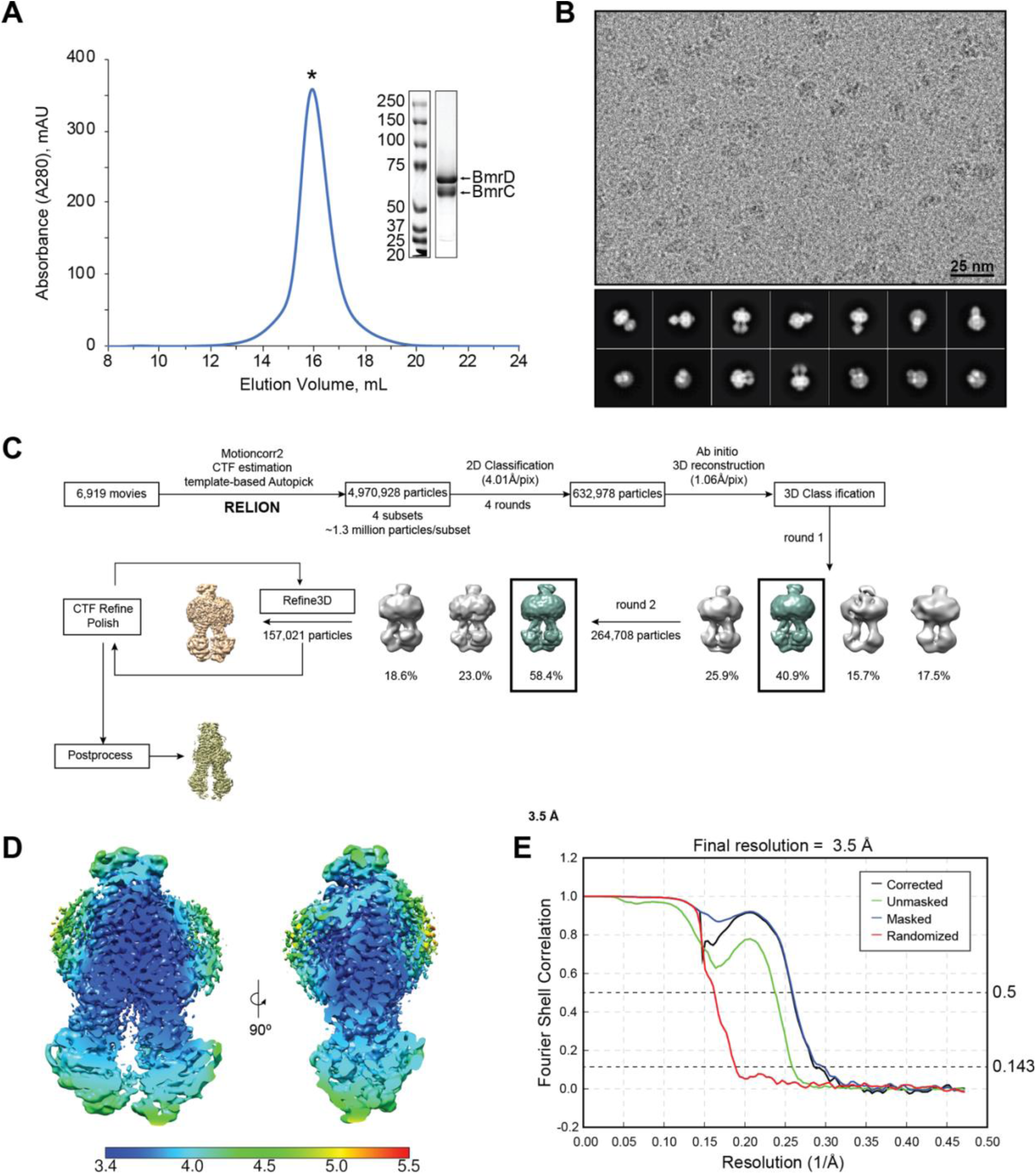
Cryo-EM data processing of BmrCD-QQ in the Hoechst-33342 and ATP-bound state. **A**) Chromatogram and SDS-PAGE analysis of the peak composition for BmrCD-QQ purified by size exclusion chromatography over a Superose 6 column equilibrated in 0.06% digitonin containing buffer prior to sample preparation for cryo-EM data collection. **B**) Representative micrograph and 2D classification results from cryo-EM data collected for BmrCD-QQ in the presence of equimolar ATP/MgCl_2_ and excess Hoechst-33342. **C**) Flowchart of data processing in RELION 3.1^33^. **D**) A slice through the EM volume colored according to local resolution. **E**) FSC curves calculated in RELION 3.1.

**Supplementary Figure 2.**
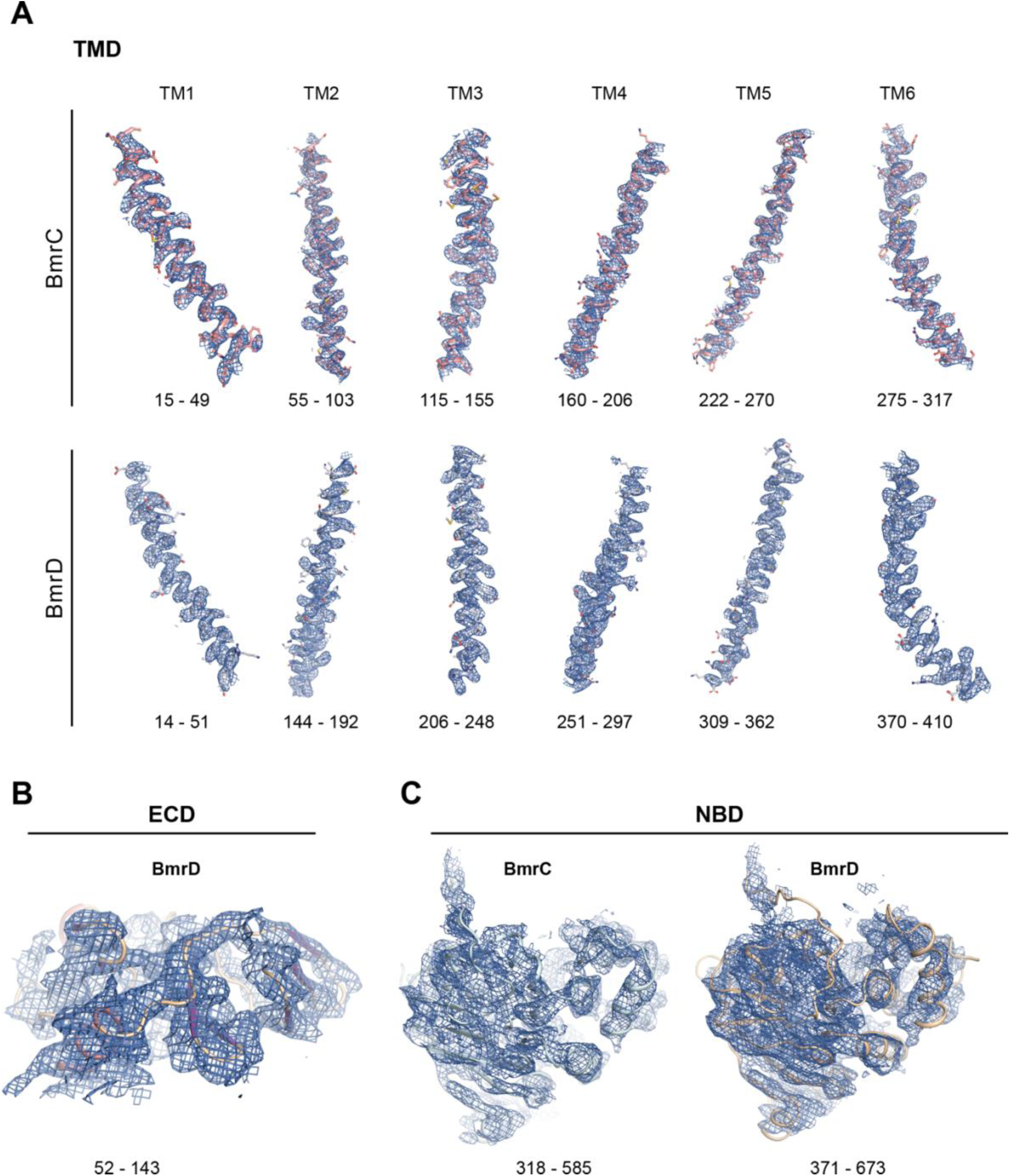
Cryo-EM map data quality. Model and corresponding density of BmrCD-QQ **A**) transmembrane helices (TM1-6), **B**) extracellular domain (ECD), and **C**) nucleotide binding domains (NBD).

**Supplementary Figure 3.**
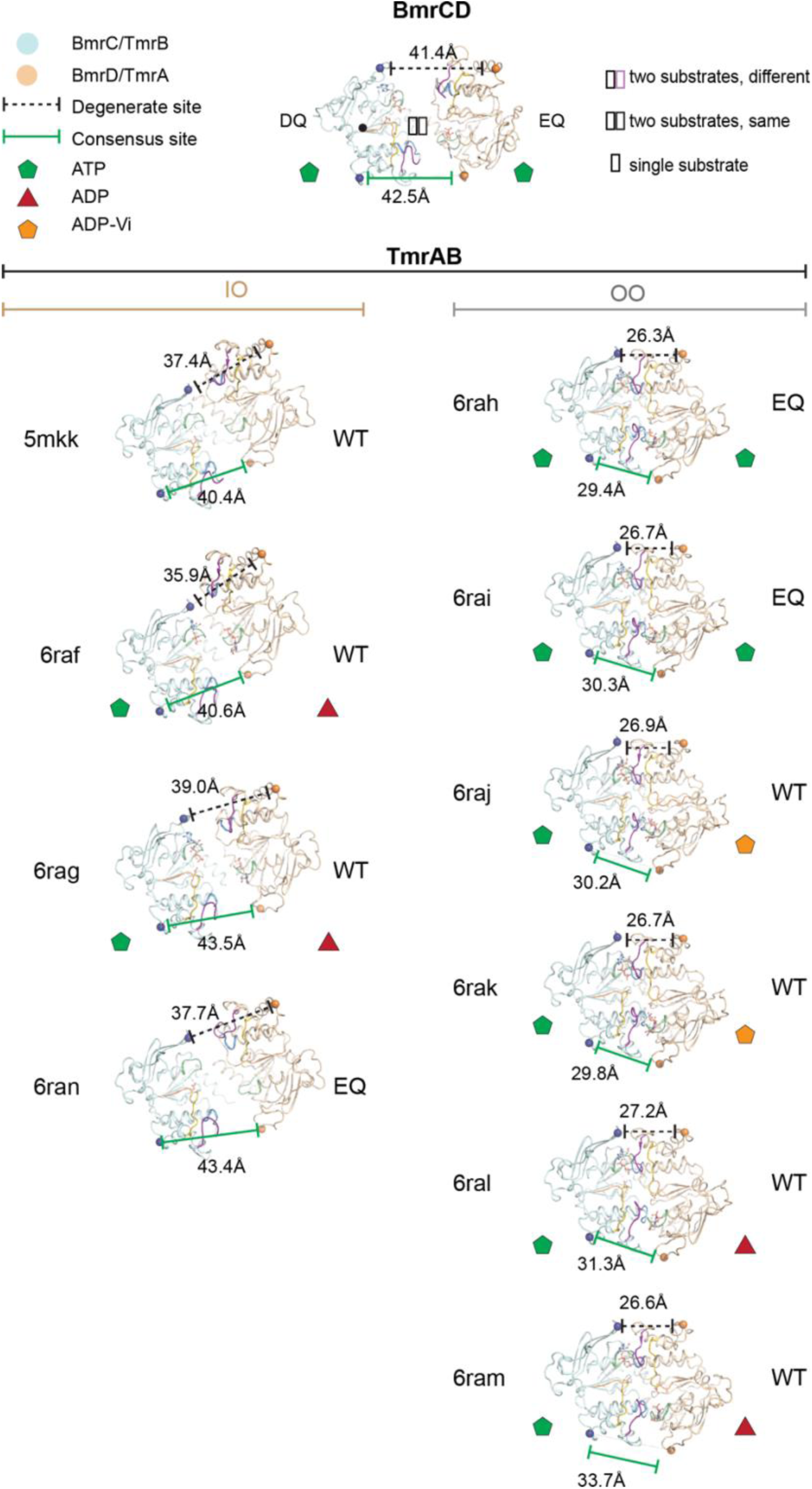
Analysis of NBD symmetry in BmrCD compared to TmrAB. Comparison of NBD geometry between BmrCD and different states of TmrAB determined by x-ray crystallography (PDB ID: 5mkk^54^) and cryo-EM^4^. The noncanonical NBD in BmrC and TmrB are colored *cyan*, whereas the consensus NBDs of BmrD and TmrA are colored *orange*. The orientations shown are relative to BmrCD and in which TmrB was aligned to BmrC (marked with a *black* dot in the center to denote the reference orientation). Structures of TmrAB are shown grouped by architectures adopting inward-facing open states (IO) or outward-facing states (OO).

**Supplementary Figure 4.**
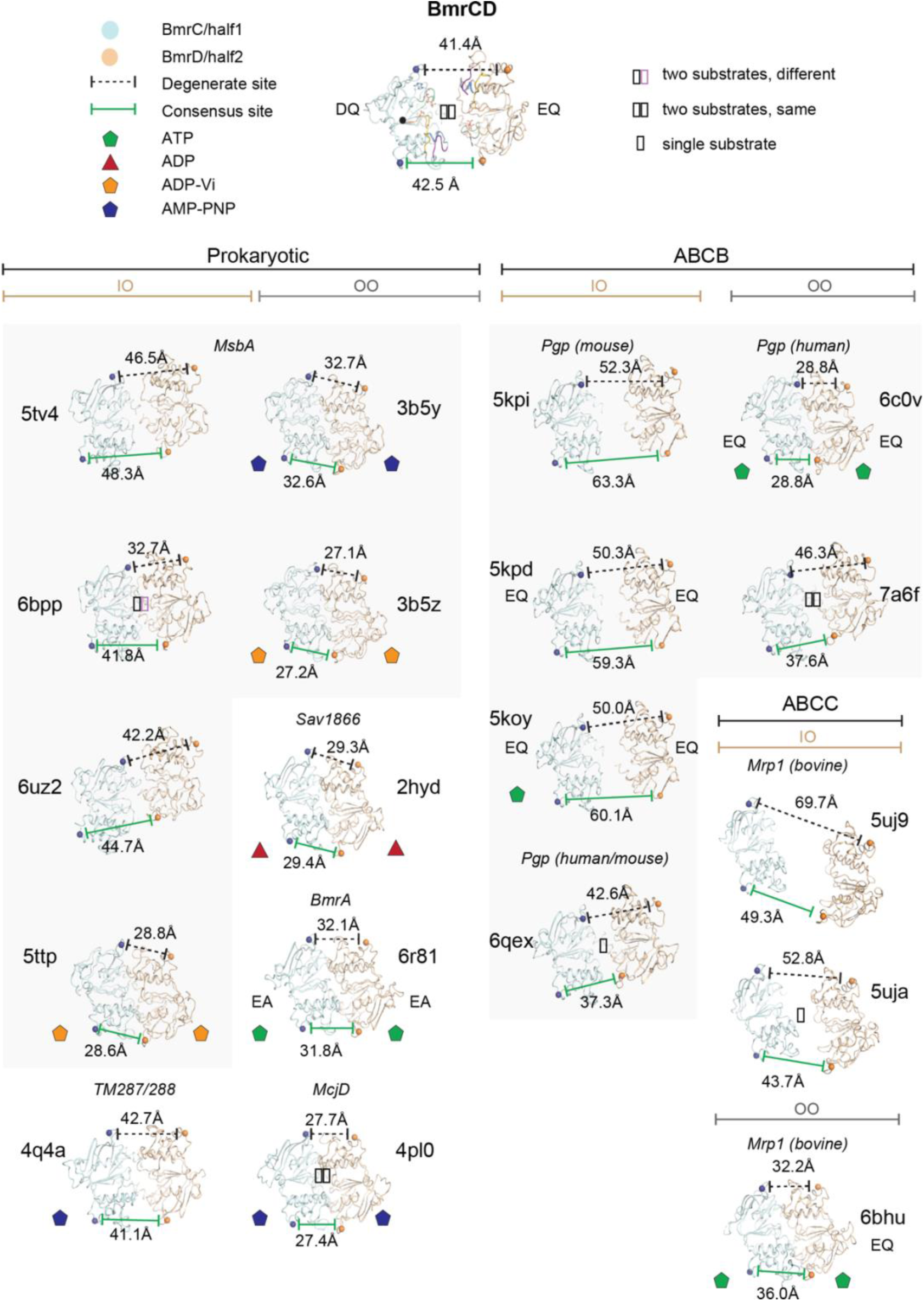
Analysis of NBD symmetry in BmrCD compared to representative ABC transporters. Comparison of NBD geometry between BmrCD and structures of related ABC transporters from prokaryotes (MsbA PDB IDs: 5tv4^18^, 6bpp^19^, 6uz2^20^, 3b5y^9^, and 3b5z^9^, TM2987/288 PDB ID: 2hyd^10^, BmrA PDB ID: 6r81 (unpublished); McjD PDB ID: 4pl0^8^), the ABCB (or Pgp) family^5, 14, 15, 22^, and ABCC family^6, 7^. The noncanonical NBDs, or the first NBD (half 1) in homomeric or single polypeptide transporters, are colored *cyan*, whereas the consensus NBDs, or the second NBD in homomeric or single polypeptide transporter, are colored *orange*. The orientations shown are relative to BmrCD in which half 1 was aligned to BmrC (marked with a black dot in the center to denote the reference orientation). Structures are shown grouped by architectures adopting inward-facing open states (IO) or outward-facing states (OO).

**Supplementary Figure 5.**
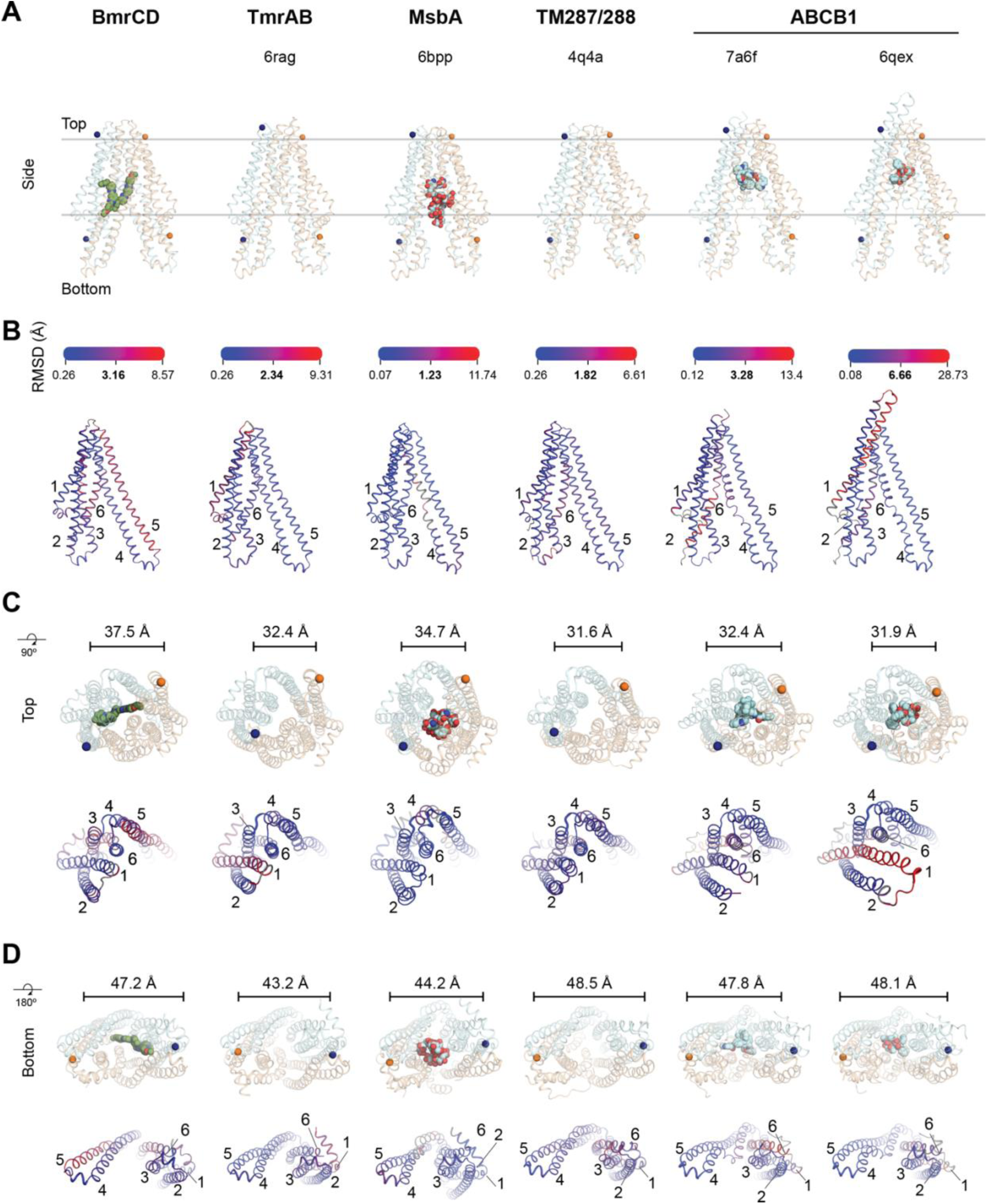
Analysis of BmrCD TMD in comparison to closely related ABC exporters. **A**) Overview of the transmembrane domain (TMD) geometry in BmrCD and related inward-facing states of related prokaryotic (PDB IDs: 6rag^4^, 6bpp^19^, and 4q4a^21^) and occluded human ABC transporters bound with inhibitor (PDB ID: 7a6f^15^) or substrate (PDB ID: 6qex^14^). The extracellular domain of BmrD and both nucleotide binding domains have been omitted for clarity. Sites on the extracellular and intracellular gates labeled for DEER experiments in BmrCD and the homologous residues in the related exporters are shown as spheres and colored the same as Figures 1 of the main text, where the non-canonical nucleotide-binding site containing half are colored *cyan* (BmrC in BmrCD), and the canonical NBS containing half are colored *orange* (BmrD in BmrCD). **B**) The Cα RMSD resulting from superposition of each TMD half within each structure is mapped onto the structure of the non-canonical half. The scale bar above represents the minimum, median, and maximum RMSD calculated for each superposition. The geometry and RMSD distribution in each are shown for the **C**) outside-facing (top) and **D**) inside-facing (bottom) regions of the TMDs to highlight the variation in asymmetry in each structure. Numbers in the RMSD representation correspond to the transmembrane helix number.

**Supplementary Figure 6.**
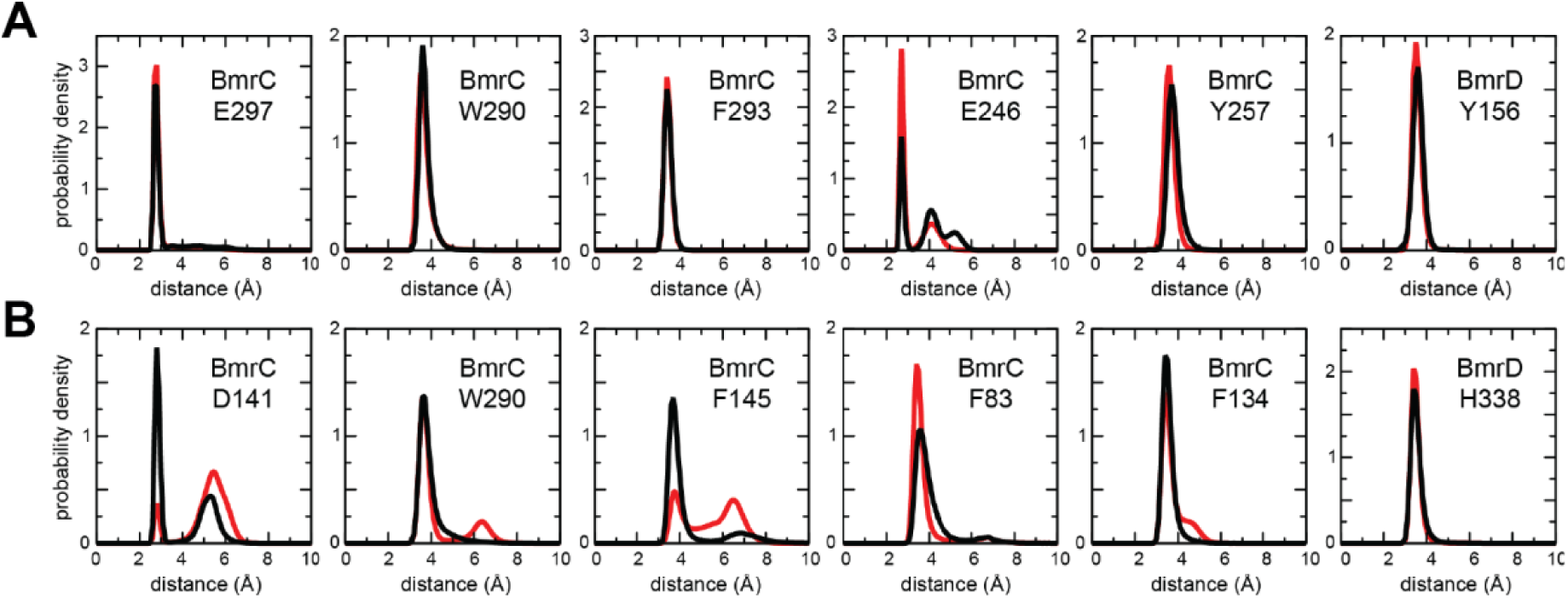
Examination of the Hoechst-33342 binding pose with MD simulations. Data is shown for two independent MD trajectories (*black*, *red*) of the BmrCD construct shown in Fig. 4E, with two Hoechst molecules bound. **A**) Quantification of ionic and aromatic interactions between Hoechst molecule ‘1’ and residues in BrmC, in terms of probability distributions of the minimum distance between the ligand and each sidechain (excluding hydrogens). The only significant contact of Hoechst molecule ‘1’ with BrmD is also indicated. **B**) Same as **A**), for molecule ‘2’.

**Supplementary Figure 7.**
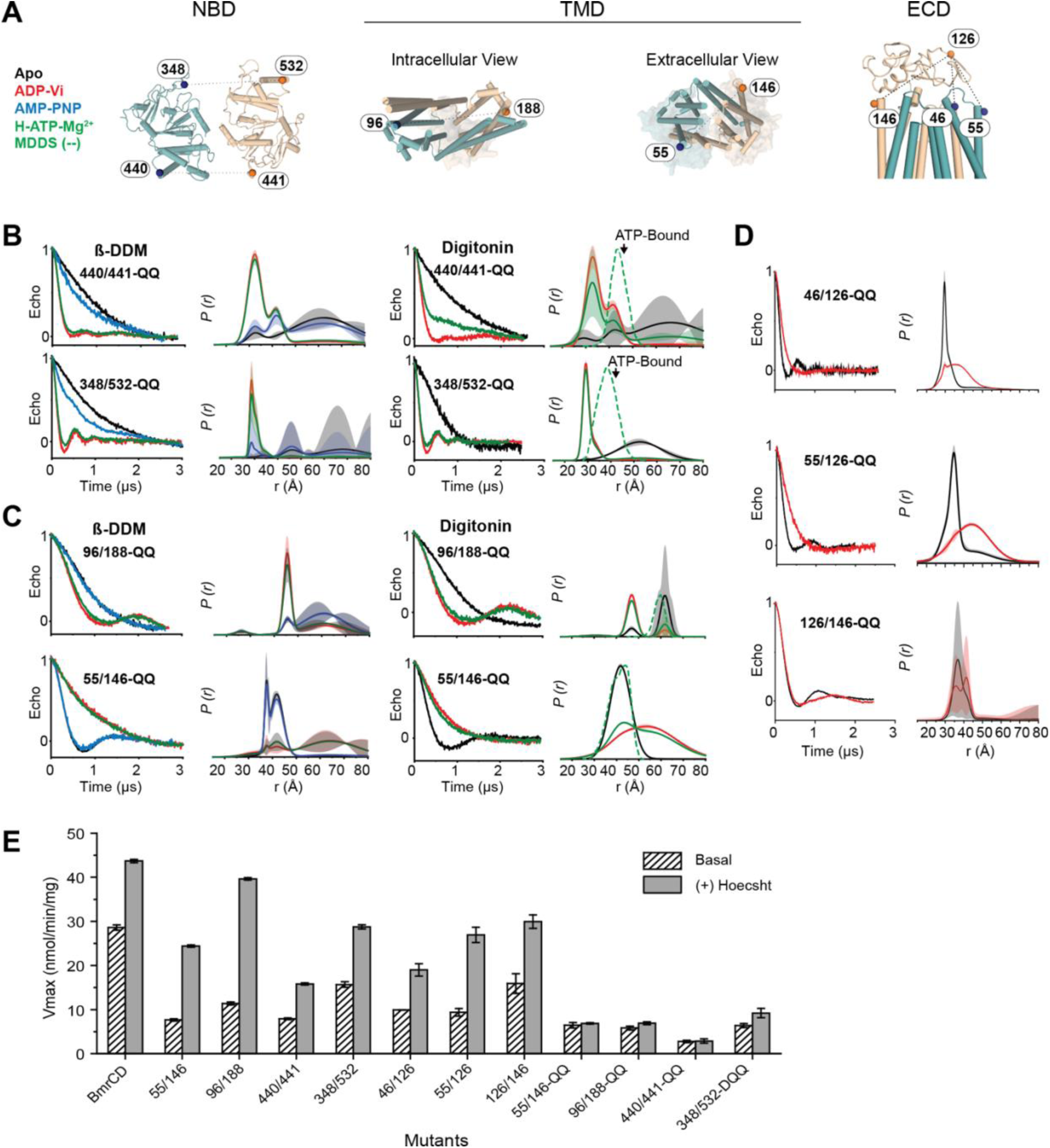
**A**) Cartoon representations of BmrCD subdomains highlighting the position of spin-labeled cysteine pairs generated in the nucleotide binding domain (NBD), transmembrane domain (TMD), and the extracellular domain (ECD). DEER decay signals and distance distributions for spin-labeled Cysteine pairs generated in the **B**) NBD, **C**) TMD, and **D**) ECD for the QQ-mutant of C-less BmrCD. The shaded regions represent confidence bands. **E**) ATP turnover rates (*V*max) in C-less BmrCD (BmrCD) and spin-labeled cysteine-pairs.

**Supplemental Table 1.** RMSD analysis of domain architecture in BmrCD

|  | RMSD (Å) |
| --- | --- |
| TMD | 3.07 |
| *TM1* | 2.2 |
| *TM2* | 1.41 |
| *TM3* | 8.58 |
| *TM4* | 4.99 |
| *TM5* | 2.07 |
| *TM6* | 1.86 |
| ICLs | 2.36 |
| *ICL1* | 1.32 |
| *ICL2* | 1.19 |
| NBD | 2.85 |

TMD: transmembrane domain; ICLs: intracellular loops; NBD: nucleotide-binding domain

**Supplemental Table 2.**
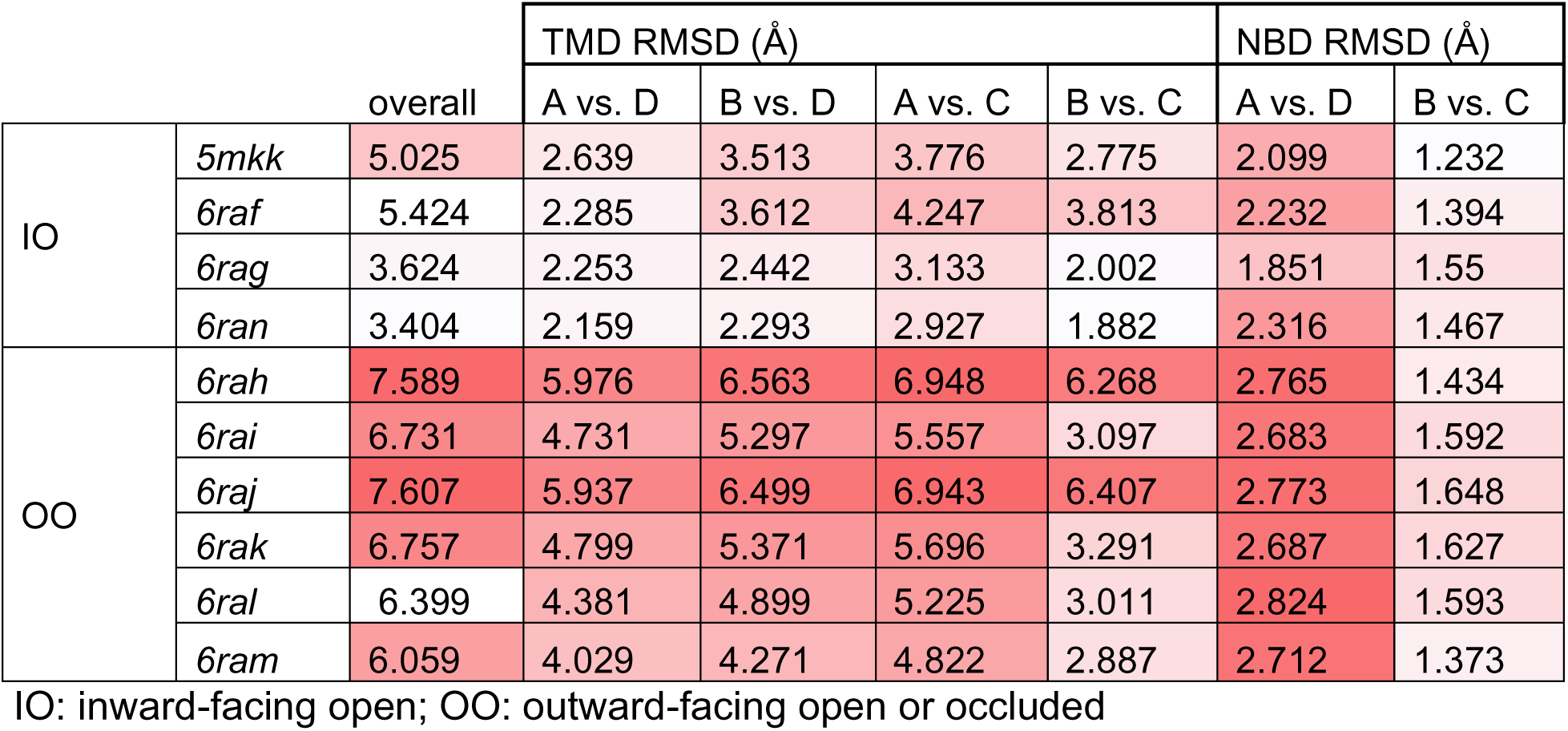
BmrCD vs. TmrAB RMSD Analysis

